# Association of simple renal cysts and chronic kidney disease with large abdominal aortic aneurysm

**DOI:** 10.1101/630558

**Authors:** Milena Miszczuk, Verena Müller, Christian E. Althoff, Andrea Stroux, Daniela Widhalm, Andy Dobberstein, Andreas Greiner, Helena Kuivaniemi, Irene Hinterseher

## Abstract

Abdominal aortic aneurysms (AAA) primarily affect elderly men who often have many other diseases, with similar risk factors and pathobiological mechanisms to AAA. The aim of this study was to assess the prevalence of simple renal cysts (SRC), chronic kidney disease (CKD), and other kidney diseases (e.g. nephrolithiasis) among patients presenting with AAA. Two groups of patients (100/group), with and without AAA, from the Surgical Clinic Charité, Berlin, Germany, were selected for the study. The control group consisted of patients who were evaluated for a kidney donation (n = 14) and patients who were evaluated for an early detection of a melanoma recurrence (n = 86). The AAA and control groups were matched for age and sex. Medical records were analyzed and computed tomography scans were reviewed for the presence of SRC and nephrolithiasis. SRC (73% vs. 57%; p<0.001) and CKD (31% vs. 8%; p<0.001) were both more common among AAA than control group patients. On multivariate analysis, CKD, but not SRC, showed a strong association with AAA. Knowledge about pathobiological mechanisms and association between CKD and AAA could provide better diagnostic and therapeutic approaches for these patients.

## Introduction

Abdominal aortic aneurysm (AAA) is the most common type of aneurysm, and is defined as an abdominal aortic diameter >3 cm [1]. According to recent literature, the prevalence of AAA has decreased in the last decades and is 1–2% [2]. This change can be primarily attributed to a decreased prevalence of smoking [2, 3]. The prevalence of AAA, however, increases with age and is 4.1%–14.2% in men and 0.35 – 6.2% in women > 65 years [4, 5].

As AAA is asymptomatic in the majority of cases [6], it is often initially detected as an incidental finding during ultrasound or computed tomography (CT) examinations. Unfortunately, many AAA cases remain undetected until rupture. The mortality rate of a ruptured AAA is estimated to be 74–90% [7, 8], with a 32–83% pre-hospital mortality rate [7, 9]. One way to reduce this trend, is to implement a national AAA screening program to detect AAA before rupture [10]. Such a program was launched successfully in the USA in 2007 [11] and in the Great Britain in 2009 [12].

A number of risk factors for AAA have been identified. The four primary risk factors are male sex, age > 65 years, smoking and a positive family history [13-18]. Several diseases appear to often co-exist with AAA, including chronic obstructive pulmonary disease (COPD) [13, 19, 20], different types of hernia [20], gallstones [21], and simple renal cysts (SRC) [22]. SRC is a common disease, with increased prevalence in older patients, affecting 24–27% of those > 50 years of age [23, 24]. In older individuals, SCRs are even more common; Carrim et al. found an overall prevalence of 41% [25], while Chang et al. reported a prevalence of 35% in >60-year olds [26].

The co-occurrence of AAA and SRC [22, 27] can be explained by shared risk factors, e.g. older age [26, 28, 29], male sex [25, 30], hypertension [31] and smoking [26]. Ito et al. [22] stated “*the presence of renal cysts shows the strongest independent association with AAA among patients belonging to the 65 to 74 years old group and over 75 years old group*”. The exact pathogenesis of SRC remains unclear, but it is intriguing that both diseases demonstrate increased matrix metalloproteinase (MMP) levels, in the aortic wall in AAA patients [32] and in the cystic fluid in patients with SRCs [33].

Given the potentially shared pathophysiology between SRC and AAA, the primary aim of this study was to assess the prevalence of SRC and other kidney diseases among AAA patients, and compare the results to a group of age- and sex-matched non-AAA patients from the same hospital.

## Materials and methods

The study was approved by the Charité Ethics Committee (approval number: EA1/309/16). Since the study was a retrospective review of medical and imaging records, no informed consent from the patients was required according to the study approval.

### Study groups

This study was a retrospective review of patients’ medical records including radiology records. Two groups of patients (100/group) were compared in the study. All 200 patients had undergone a computed tomography-angiography (CTA) scan. The first group included patients, who underwent AAA surgery in 2004 – 2012 at Charité Clinic Campus Mitte in Berlin, Germany. Surgeries were performed either as elective (unruptured AAA; n=94) or as emergency (ruptured AAA; n= 6) operations. The exclusion criteria were an abdominal aortic diameter <3 cm, AAA operation before 2004, diagnosis of rare genetic disorder such as Marfan syndrome or Ehlers-Danlos syndrome, and the presence of any other arterial aneurysm. One AAA patient was diagnosed with Marfan syndrome and was, therefore, excluded from further analyses, leaving 99 AAA patients for the analyses presented here. For the AAA patients, the pre-operative scans were used.

The control group (n=100) included patients without AAA investigated at the Institute of Radiology of Charité Clinic, Berlin, Germany, and consisted of patients who were evaluated for a kidney donation in 2005 – 2014 (n = 14) and patients who were evaluated for an early detection of a melanoma recurrence (n = 86). We chose this group of patients as a control group for the following reasons: 1) they were also examined by abdominal CTA; 2) melanoma is an age-related disease and a disease of a different organ, not the aorta; and 3) they were from the same hospital system. Also, there were no differences in the mean height, weight, or BMI between the AAA and control groups [34].

AAA patients and controls were matched on sex and age (± 2 years). For the AAA patients, age at the time of the first AAA diagnosis was used for this analysis. If this information was missing, age at the time of AAA surgery was taken. For the control group, age during the CTA scan was used for the analysis.

### Clinical data

For the analysis of the CTA scans, Centricity eRadCockpit Software (GE Healthcare, Chalfont St Giles, Great Britain) was used. First, written reports from board-certified radiologists were reviewed by one of the authors (M.M.). As SRCs are common, sometimes they were not described as a diagnosis in the report. For that reason, the CTA scans were assessed again for the presence of SRC (Fig 1) and kidney stones. Results were discussed with a board-certified radiologist (C.E.A.).

**Fig. 1.**
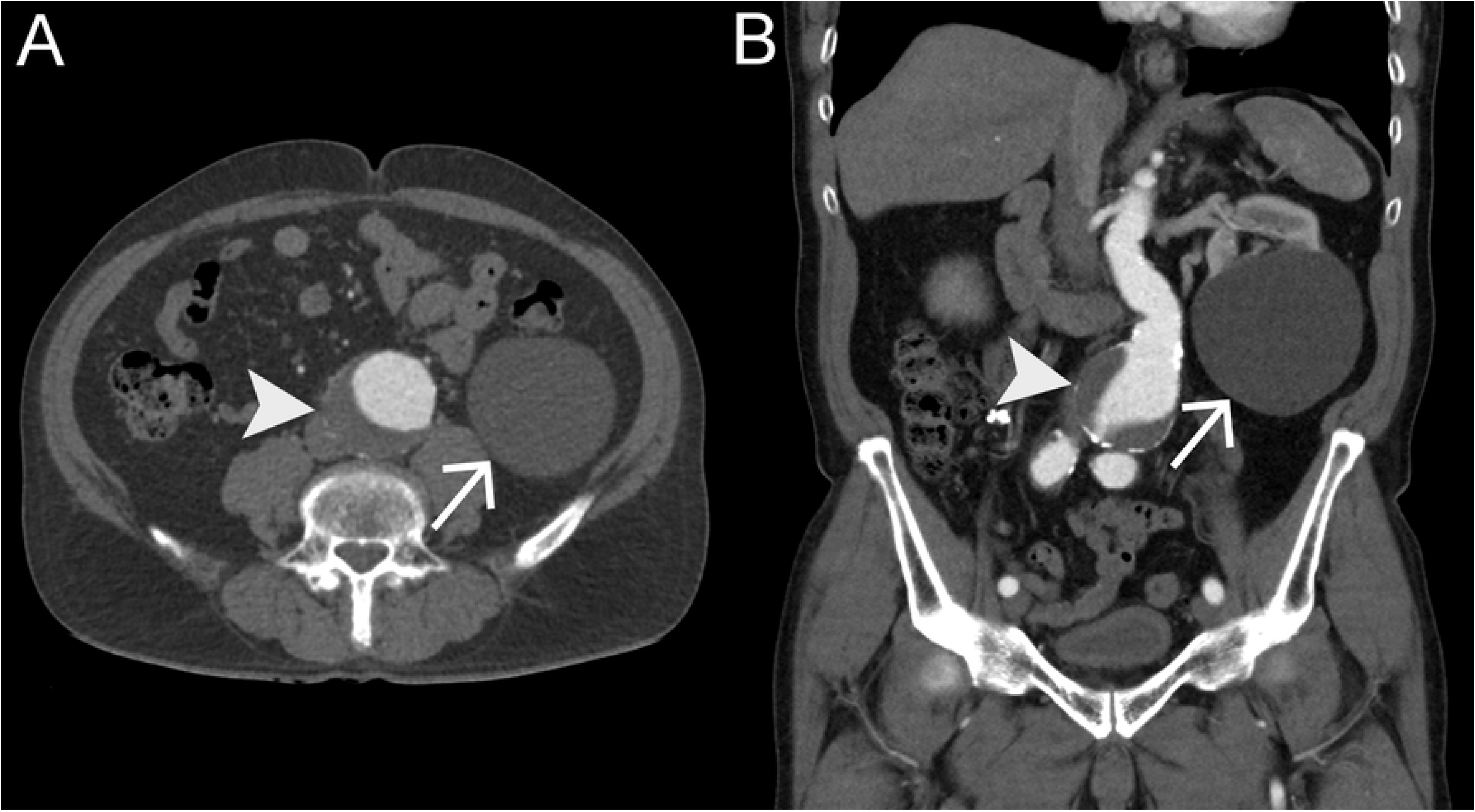
Simple renal cyst detected in a CT scan. Contrast-enhanced CT scan of the abdomen in axial (A) and coronal (B) plane, arterial phase, demonstrates an AAA (arrow head) with mural thrombus and patent lumen and a large hypodense mass on the lower left renal pole, representing a simple renal cyst (arrow).

Individual data on all study patients for all variables used in the study are available in the Supplementary file.

SRC (ICD-10: N28.1) were divided into subgroups according to their size: Group 1: ≤1 cm; Group 2: 1.01 – 3.0 cm; Group 3: 3.1 – 5.0 cm; and Group 4: >5 cm. SRCs were also classified using Bosniak Classification System: I: simple, benign cysts; II: minimally complicated benign cystic lesions; III: more complicated cystic lesions; and IV: cystic carcinoma [35].

Nephrolithiasis (ICD-10: N20.0) was defined as a presence of kidney stones on the CTA scans.

Additional patient data [34] were collected from the medical records using the patient data management program SAP (SAP SE, Walldorf, Germany). Information about the presence of chronic kidney disease (CKD) was collected. CKD was defined as a presence of the following ICD-10 diagnostic codes in the medical records: N18.1 for CKD stage 1, N18.2 for CKD stage 2, N18.3 for CKD stage 3, N18.4 for CKD stage 4, N18.5 for CKD stage 5, and Z94.0 for renal transplantation. Additional diseases in the same study groups are described in another study [34].

### Statistical analysis

For statistical analyses SPSS Statistics Version 22 for Windows (IBM, Armonk, New York, USA) was used. First, a univariable analysis was carried out. For quantitative variables, the mean, median, standard deviation, minimal and maximal values were determined. The categorical variables were analyzed using cross-tabulation. The differences between the two groups were determined using Mann-Whitney U test or chi-squared test (Fisher’s exact test) where appropriate. A difference was defined significant, if p≤0.05.

The univariable analysis was followed by a multivariable analysis to identify independent risk factors. Significant values from the univariable analysis were included in a multiple logistic regression model. This included the following parameters: ever smoker, peripheral artery disease (PAD), pack years of smoking, incisional hernia, any hernia, congestive heart failure, American Society of Anesthesiologists (ASA) score, diabetes mellitus, coronary bypass, creatinine, COPD, current smoker, coronary artery disease, diverticulosis, platelet count [34] and SRC. Parameters with >50% of the values missing were excluded from the analysis. A forward and backward analysis was performed. Odds ratios (OR) and a 95% confidence intervals (CI) were calculated.

## Results

Our study included 99 (78 male and 21 female) AAA patients and 100 (79 male and 21 female) age- and sex-matched controls. Altogether, 30.3% AAA and 8% control patients had CKD diagnosis in their medical records (p<0.001; Table 1). The distribution of CKD stages was also statistically significantly different between the study groups (p=0.002; Table 1). One AAA patient received a renal transplantation due to the CKD. Nephrolithiasis was found in 2% of the AAA and 7.1% of the control patients (p=0.1). SRCs were found amongst 72.7% AAA patients and 57% controls, resulting in a statistically significant difference (Table 1; p=0.009). Among AAA patients, two had been diagnosed with autosomal dominant polycystic kidney disease (ADPKD) and were excluded from further evaluations.

**Table 1.** Comparison of simple renal cysts and chronic kidney disease between study groups.

| Variable | AAA group |  | Control group |  | *p*<sup>a</sup> |
| --- | --- | --- | --- | --- | --- |
|  | *With variable*<br>*n* | *Data available*<br>*n* | *With variable*<br>*n* | *Data available*<br>*n* |  |
| CKD, all stages<sup>b</sup> | 30 | 99 | 8 | 100 | <0.001 |
| CKD stage 1/2/3/4/5/RTX<sup>c</sup> | 5/9/11/1/3/1 | 99 | 1/4/3/0/0/0 | 100 | 0.002 |
| Nephrolithiasis | 2 | 99 | 7 | 98 | 0.10 |
| SRC (both right and left kidney)<sup>d</sup> | 72 | 99 | 57 | 100 | 0.01 |
| SRC, right kidney only | 58 | 99 | 46 | 100 | 0.04 |
| 1 – 4 SRC in right kidney<sup>d</sup> | 46 | 99 | 41 | 100 | 0.05 |
| >5 SRC in right kidney<sup>d</sup> | 12 | 99 | 5 | 100 | 0.05 |
| SRC, left kidney only<sup>d</sup> | 60 | 98 | 33 | 100 | <0.001 |
| 1 – 4 SRC in left kidney<sup>d</sup> | 48 | 98 | 29 | 100 | <0.001 |
| > 5 SRC in left kidney<sup>d</sup> | 12 | 98 | 4 | 100 | <0.001 |

<sup>a</sup>Chi-square test
<sup>b</sup>Defined as presence of ICD-10 code in medical records: N18.1 for CKD stage 1, N18.2 for CKD stage 2, N18.3 for CKD stage 3, N18.4 for CKD stage 4, N18.5 for CKD stage 5 or Z94.0 for renal transplantation.
<sup>c</sup>Defined as presence of kidney stones (ICD-10: N20.0) on the CTA scans.
<sup>d</sup>Defined as presence of simple renal cysts (ICD-10: Q61.9) on the CTA scans.
CKD: chronic kidney disease; SRC: simple renal cyst; RTX: renal transplantation.

In the right kidney, SRCs were found in 58.6% of AAA patients and in 46% of controls (p=0.039). In the AAA group, 46.5% of patients had 1–4 SRCs, and 12.1% of patients had ≥5 SRCs. In the control group, these numbers were 41% and 5%, respectively (p=0.048, for both comparisons). In the right kidney, SRCs ≤1 cm were significantly more common among AAA than control patients (p<0.001; Table 2).

**Table 2.** Simple renal cysts in the right kidney classified according to their size.

| SRC<br>classification<br>based on size | AAA group |  |  | Control group |  |  | *p*<sup>a</sup> |
| --- | --- | --- | --- | --- | --- | --- | --- |
|  | *Number of* | *Median* | *Data* | *Number of* | *Median* | *Data* |  |
|  | *SRC per* | *number of availa* |  | *SRC per* | *number of availa* |  |  |
|  | *patient* | *SRC per* | *ble* | *patient* | *SRC per* | *ble* |  |
|  | *Mean+/-SD* | *patient* | *n* | *Mean+/-SD* | *patient* | *n* |  |
| ≤1 cm | 1.15 ± 1.86 | 0 | 97 | 0.50 ± 1.45 | 0 | 100 | <0.001 |
| 1.1 cm – 3.0 cm | 0.43 ± 0.95 | 0 | 97 | 0.41 ± 0.79 | 0 | 100 | 0.99 |
| 3.1 cm – 5.0 cm | 0.10 ± 0.34 | 0 | 97 | 0.05 ± 0.26 | 0 | 100 | 0.15 |
| >5 cm | 0.02 ± 0.14 | 0 | 97 | 0.04 ± 0.20 | 0 | 100 | 0.68 |

<sup>a</sup>Mann-Whitney U test
SRC, simple renal cyst

In the left kidney, SRCs were found in 61.2% of AAA and 33% of control patients (p<0.001). In the AAA group, 49% of patients had 1–4 SRCs, and 12.2% of patients had ≥5 SRCs. In the control group, these numbers were 29% and 4%, respectively (p<0.001, for both variables). In the left kidney, both small (≤1 cm; p<0.001) and medium size (1 – 3 cm; p=0.020) SRCs were significantly more common among AAA than control patients (Table 3). In the AAA group, we also found two patients with SRCs classified as Bosniak II and two patients with SCRs classified as Bosniak III. In the control group, one patient each with Bosniak II and Bosniak III were found. There were 46 (46.5%) AAA and 22 (22%) control group patients who had bilateral SRC disease, whereas 25 (25.3%) AAA and 34 (34%) control patients had SRCs in only the left or right kidney (p=0.001).

**Table 3.** Simple renal cysts in the left kidney classified according to their size.

| SRC classification based on size | AAA | | | Controls | | | $p^a$ |
| --- | --- | --- | --- | --- | --- | --- | --- |
|  | Number of SRC per patient | Median number of SRC per | Data available, | Number of SRC per patient | Median number of SRC per | Data available, |  |

|  | *Mean+/-SD* | *patient* | *n* | *Mean+/-SD* | *patient* | *n* |  |
| --- | --- | --- | --- | --- | --- | --- | --- |
| ≤1 cm | 1.28 ± 2.08 | 1 | 96 | 0.37 ±1.17 | 0 | 100 | <0.001 |
| 1.1 cm – 3.0 cm | 0.56 ± 0.95 | 0 | 96 | 0.30 ± 0.73 | 0 | 100 | 0.02 |
| 3.1 cm – 5.0 cm | 0.10 ± 0.31 | 0 | 96 | 0.04 ± 0.20 | 0 | 100 | 0.10 |
| >5 cm | 0.01 ± 0.10 | 0 | 96 | 0.00 ± 0.00 | 0 | 100 | 0.49 |

<sup>a</sup>Mann-Whitney U test
SRC, simple renal cyst

In multivariable analyses, which included ever smoker, PAD, pack years, incisional hernia, any hernia, congestive heart failure, ASA score, diabetes mellitus, coronary bypass, creatinine, COPD, current smoker, coronary artery disease, diverticulosis, platelet count and SRC, we found a strong independent association between AAA and CKD (OR = 5.655; 95%CI = 1.785– 17.921; p=0.003). We found no direct association between SRC and AAA (OR = 1.711; 95%CI = 0.625–4.682; p=0.296), when adjusting for other variables.

## DISCUSSION

The main findings of our study were that CKD was more frequent in AAA than age- and sex-matched control patients and showed a strong association with AAA in multivariable analysis, which included ever smoker, PAD, pack years, incisional hernia, any hernia, congestive heart failure, ASA score, diabetes mellitus, coronary bypass, creatinine, COPD, current smoker, coronary artery disease, diverticulosis, platelet count and SRC. AAA patients also had a higher rate of SRC, but SRCs were not independently associated with AAA.

Our study demonstrated a strong association between AAA and CKD with 30.3% of the AAA and only 8% of the age- and sex-matched control patients diagnosed with CKD. The AAA patients also exhibited a more advanced CKD stage. Previously published studies reported a wide range of CKD prevalence (3–65%) among AAA patients [20, 36-39]. Alnassar et al. [36] and Pitoulias et al. [40] found no significant difference in the prevalence of CKD between AAA and PAD patients [20, 36]. However, patients with a large AAA (>5.5 cm) had a significantly higher rate of CKD than patients with a small AAA (13% vs. 2%) [20]. Approximately half (54.3%) of the AAA patients in our study had a large AAA (>5.5 cm) and all were operated on for AAA, and the rate of CKD was over four times as high as the rate in the study of Pitoulias et al [20]. Furthermore, similar to the findings by Chun et al. [38] and Takeuchi et al. [39], our study demonstrated an independent association of AAA and CKD in multivariable analysis, not seen in the study by Pitoulias et al. [20].

We estimated the prevalence of SRC to be 72.7% in the AAA and 57% in the control group. Based on previous literature, the SRC prevalence in general population varies 4.2–41% [25, 28], which is lower than in the current study (Table 4). Similarly, the SRC prevalence among AAA patients in the current study was higher than in most of the previous studies, which reported a prevalence of 38–69% among AAA patients and 18–45% for controls [20, 22, 27, 41-43]. A recent study by Brownstein et al. [43] analyzed a total of 35,498 patients who underwent both chest and abdominal CT imaging during a 4-year period. Altogether 18% of these patients had SRC and 2.6% had AAA. Compared with the matched population without SRC, patients with SRC demonstrated an increased prevalence of AAA (8% vs. 3%). They were also more likely to have thoracic, ascending and descending aortic aneurysms or dissections [43]. Five previous studies found an independent association between AAA and SRC in a multivariable analysis [20, 22, 27, 41, 43], but we could not confirm this in our study. However, three of those studies [22, 41, 43] examined a significantly larger patient group.

**Table 4.**
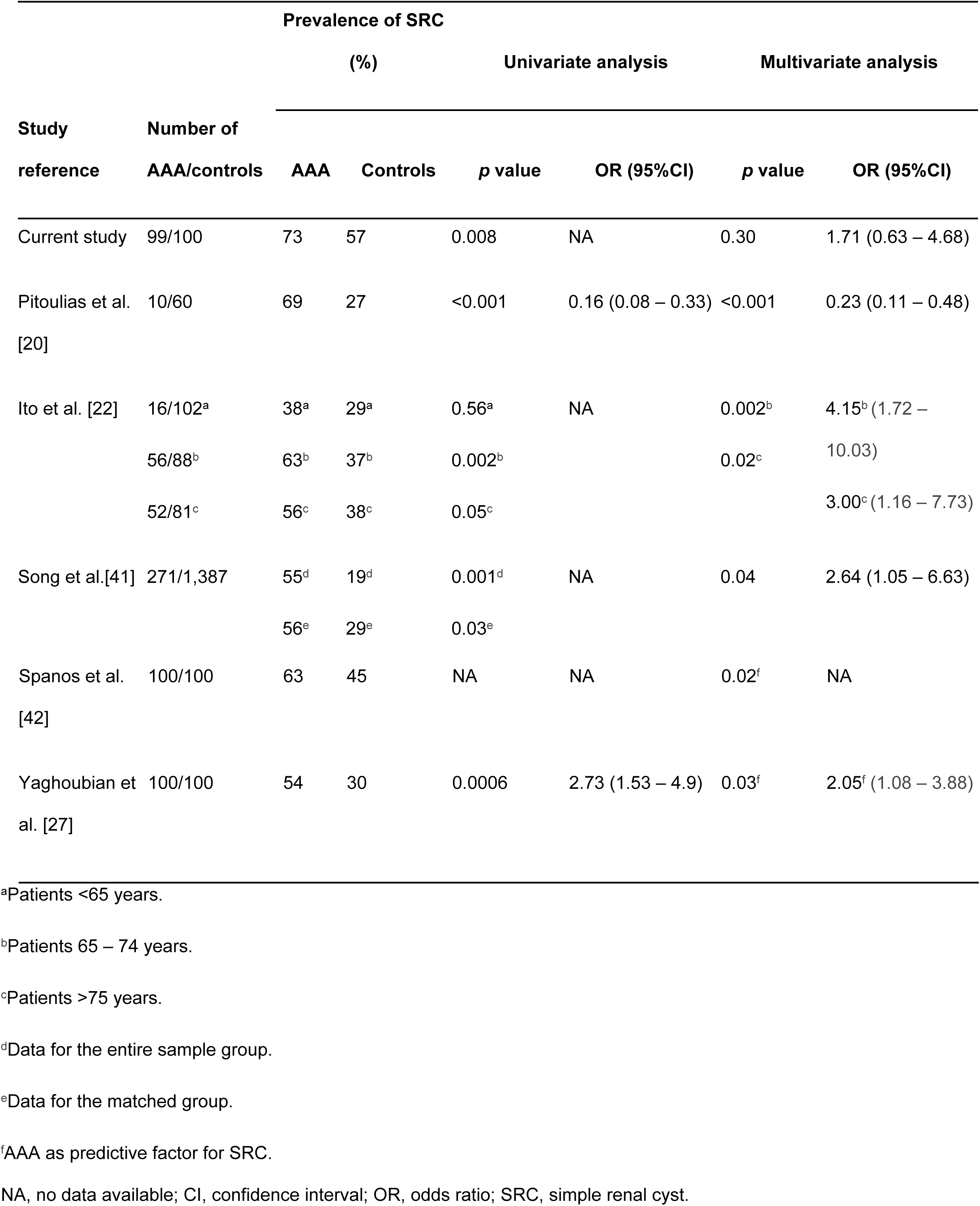
Prevalence of SRC in AAA and control patients in published studies.

An important difference between our study and the previous studies on SRCs is that in our study patients were not stratified according to age, which could explain our negative results in the multivariable analysis. Ito et al. [22] found an association between AAA and SRC in their multivariable analysis, but only in patients >65 years. Another important factor is the age of patients with and without SRC. Prior studies have confirmed that SRC develops mostly at older age [24, 25, 28, 29, 41, 42], and Ito et al. [22] and Yaghoubian et al. [27] found a significant difference in the age of patients with and without SRC.

Our study showed an asymmetric distribution of SRCs between the left and right kidney. In controls, the right kidney was more often affected than the left kidney (46% vs. 33%,), whereas in the AAA group, both kidneys were affected as often (58.6% vs. 61.2%). One previous study demonstrated that a bilateral appearance of SRC increased the risk of hypertension (OR = 3.48, 95%CI = 2.12–5.71) [44]. Also the presence of multiple SRC (≥ 2) was associated with an increased risk of hypertension [44]. Previous studies have also examined the correlation between SRC size and hypertension, but an association with hypertension was found only for SRCs > 1 cm in diameter [44].

SRC and AAA share some common risk factors, e.g. older age, male sex and hypertension [25, 26, 28, 29, 31, 45], and some studies also mention smoking as a possible risk factor for SRC [26]. Molecular studies suggest that MMPs play a role in the pathophysiology of SRC and AAA [33]. Furthermore, one study on 108 autopsies showed a correlation between the diameter of the abdominal aorta and the number of SRC [46].

Further research on kidney diseases and AAA is not only of academic, but also of clinical interest. Nowadays, the majority of AAA are repaired using endovascular aneurysm repair (EVAR), which requires a contrast agent administration, known to be nephrotoxic. Only one study has investigated the kidney function in patients with SRC after EVAR [42], and found that patients with SRC had slightly higher creatinine levels, both before and after surgery, but the difference was not statistically significant [42]. There was no significant difference in the creatinine levels after EVAR [42], leading to the conclusion that kidney function is not affected by the presence of SRC. As we have reported previously, in the current study population, the AAA group patients had significantly higher creatinine levels than the control group patients [34].

Nephrolithiasis is a common problem in the elderly population. A higher prevalence of nephrolithiasis has been reported in patients with SRC [26] and ADPKD [47]. The relationship between renal stones and AAA has not been investigated previously. As SRC and ADPKD appear frequently in AAA patients [22, 27], one might expect that nephrolithiasis affects AAA patients as well. In our study population, we found no association between AAA and nephrolithiasis. Nonetheless, when examining a patient with a renal colic, one should consider a symptomatic or ruptured AAA as a potential differential diagnosis.

The association between AAA and CKD also requires further research by examining the role CKD plays on the development and progression of AAA. It has been reported that blood vessel walls in patients with CKD are thinner, which can increase the risk for rupture [48]. The co-occurrence of AAA and CKD also has several clinical implications, since CKD increases the rate of complications after surgery [49]. The clot inside the aneurysm sac can also impair the blood perfusion of renal arteries (e.g. by embolization).

The main limitation of our study is the fact that this was a retrospective single-center study with a small number of patients and controls. Another possible limitation is the fact that the control group consisted of patients with melanoma and those evaluated for kidney donation and their comorbidity profile might not be representative of the general population. Also, our study evaluated patients with larger AAAs which were treated surgically. This might have caused a selection bias and the results might not be representative of all AAA patients. A major advantage of our study was matching of patients and controls on sex and age minimizing the confounding effects attributed to these factors.

A better understanding about the pathophysiology of AAA will facilitate the development of pharmacotherapies for AAA. Also, this knowledge could be used for a better risk stratification. By introducing a national screening program in every country, AAA could be detected earlier. It is also important to consider the possible complications arising from CKD, both after open aneurysm repair, and EVAR. This group of patients should be given special attention and risk factor analysis should be carried out. The risk of rupture should be high enough to justify the risk of surgery. Further research is also needed on patients with small AAA who develop problems in kidney function. It remains to be determined if renal function is also affected by an expansion and different types of AAA where the renal arteries are involved.

## Funding

This research received no specific grant from any funding agency in the public, commercial, or not-for-profit sectors.

## Data Availability Statement

All relevant data are within the paper and the files part of the Supporting Information.

## Supporting information

S1 File. Dataset containing all information on every patient included in the study (Excel format).

## References

[1] Moll FL, Powell JT, Fraedrich G, Verzini F, Haulon S, Waltham M, et al. Management of abdominal aortic aneurysms clinical practice guidelines of the European society for vascular surgery. Eur J Vasc Endovasc Surg 2011;41 Suppl 1:S1–S58.

[2] Lilja F, Wanhainen A, Mani K. Changes in abdominal aortic aneurysm epidemiology. J Cardiovasc Surg (Torino) 2017;58:848–53.

[3] Lederle FA. The rise and fall of abdominal aortic aneurysm. Circulation 2011;124:1097–9.

[4] Cornuz J, Sidoti Pinto C, Tevaearai H, Egger M. Risk factors for asymptomatic abdominal aortic aneurysm: systematic review and meta-analysis of population-based screening studies. Eur J Public Health 2004:343–9.

[5] Fleming C, Whitlock EP, Beil TL, Lederle FA. Screening for abdominal aortic aneurysm: a best-evidence systematic review for the U.S. Preventive Services Task Force. Ann Intern Med 2005;142:203–11.

[6] Keisler B, Carter C. Abdominal aortic aneurysm. Am Fam Physician 2015;91:538–43.

[7] Reimerink JJ, van der Laan MJ, Koelemay MJ, Balm R, Legemate DA. Systematic review and meta-analysis of population-based mortality from ruptured abdominal aortic aneurysm. Br J Surg 2013;100:1405–13.

[8] Acosta S, Ogren M, Bengtsson H, Bergqvist D, Lindblad B, Zdanowski Z. Increasing incidence of ruptured abdominal aortic aneurysm: a population-based study. J Vasc Surg 2006;44:237–43.

[9] Guirguis-Blake JM, Beil TL, Sun X, Senger CA, Whitlock EP. Primary care screening for abdominal aortic aneurysm: an evidence update for the U.S. preventive services task force. Evidence Synthesis No 109 AHRQ Publication No 14-05202-EF-1 Rockville, MD: Agency for Healthcare Research and Quality 2014.

[10] Siso-Almirall A, Kostov B, Navarro Gonzalez M, Cararach Salami D, Perez Jimenez A, Gilabert Sole R, et al. Abdominal aortic aneurysm screening program using hand-held ultrasound in primary healthcare. PLoS One 2017;12:e0176877.

[11] Lederle FA. Screening for AAA in the USA. Scand J Surg 2008;97:139–41.

[12] Benson RA, Poole R, Murray S, Moxey P, Loftus IM. Screening results from a large United Kingdom abdominal aortic aneurysm screening center in the context of optimizing United Kingdom National Abdominal Aortic Aneurysm Screening Programme protocols. J Vasc Surg 2016;63:301–4.

[13] Simoni G, Pastorino C, Perrone R, Ardia A, Gianrossi R, Decian F, et al. Screening for abdominal aortic aneurysms and associated risk factors in a general population. Eur J Vasc Endovasc Surg 1995;10:207–10.

[14] Alcorn HG, Wolfson SK, Jr., Sutton-Tyrrell K, Kuller LH, O’Leary D. Risk factors for abdominal aortic aneurysms in older adults enrolled in The Cardiovascular Health Study. Arterioscler Thromb Vasc Biol 1996;16:963–70.

[15] Blanchard JF, Armenian HK, Friesen PP. Risk factors for abdominal aortic aneurysm: results of a case-control study. Am J Epidemiol 2000;151:575–83.

[16] Lederle FA, Johnson GA, Wilson SE, Chute EP, Hye RJ, Makaroun MS, et al. The aneurysm detection and management study screening program. Arch Intern Med 2000;160:1425–30.

[17] Kent KC, Zwolak RM, Egorova NN, Riles TS, Manganaro A, Moskowitz AJ, et al. Analysis of risk factors for abdominal aortic aneurysm in a cohort of more than 3 million individuals. J Vasc Surg 2010;52:539–48.

[18] Wahlgren CM, Larsson E, Magnusson PK, Hultgren R, Swedenborg J. Genetic and environmental contributions to abdominal aortic aneurysm development in a twin population. J Vasc Surg 2010;51:3–7.

[19] Meijer CA, Kokje VB, van Tongeren RB, Hamming JF, van Bockel JH, Moller GM, et al. An association between chronic obstructive pulmonary disease and abdominal aortic aneurysm beyond smoking: results from a case-control study. Eur J Vasc Endovasc Surg 2012;44:153–7.

[20] Pitoulias GA, Donas KP, Chatzimavroudis G, Torsello G, Papadimitriou DK. The role of simple renal cysts, abdominal wall hernia, and chronic obstructive pulmonary disease as predictive factors for aortoiliac aneurysmatic disease. World J Surg 2012;36:1953–7.

[21] Schuster JJ, Raptopoulos, V., Baker, S.P. Increased prevalence of cholelithiasis in patients with abdominal aortic aneurysm: sonographic evaluation. Am J Roentgenol 1989;152:509–11.

[22] Ito T, Kawaharada N, Kurimoto Y, Watanabe A, Tachibana K, Harada R, et al. Renal cysts as strongest association with abdominal aortic aneurysm in elderly. Ann Vasc Dis 2010;3:111–6.

[23] Laucks SP, Jr,, McLachlan MS. Aging and simple cysts of the kidney. Br J Radiol 1981;54:12–4.

[24] Tada S, Yamagishi J, Kobayashi H, Hata Y, Kobari T. The incidence of simple renal cyst by computed tomography. Clin Radiol 1983;34:437–9.

[25] Carrim ZI, Murchison JT. The prevalence of simple renal and hepatic cysts detected by spiral computed tomography. Clin Radiol 2003;58:626–9.

[26] Chang CC, Kuo JY, Chan WL, Chen KK, Chang LS. Prevalence and clinical characteristics of simple renal cyst. J Chin Med Assoc 2007;70:486–91.

[27] Yaghoubian A, de Virgilio C, White RA, Sarkisyan G. Increased incidence of renal cysts in patients with abdominal aortic aneurysms: a common pathogenesis? Ann Vasc Surg 2006;20:787–91.

[28] Chin HJ, Ro H, Lee HJ, Na KY, Chae DW. The clinical significances of simple renal cyst: Is it related to hypertension or renal dysfunction? Kidney Int 2006;70:1468–73.

[29] Mosharafa AA. Prevalence of renal cysts in a Middle-Eastern population: an evaluation of characteristics and risk factors. BJU Int 2008;101:736–8.

[30] Terada N, Ichioka K, Matsuta Y, Okubo K, Yoshimura K, Arai Y. The natural history of simple renal cysts. J Urol 2002;167:21–3.

[31] Pedersen JF, Emamian SA, Nielsen M, B. Significant association between simple renal cysts and arterial blood pressure. Br J Ur 1997;79:688–92.

[32] Liapis CD, Paraskevas KI. The pivotal role of matrix metalloproteinases in the development of human abdominal aortic aneurysms. Vasc Med 2003;8:267–71.

[33] Harada H, Furuya M, Ishikura H, Shindo J, Koyanagi T, Yoshiki T. Expression of matrix metalloproteinase in the fluids of renal cystic lesions. J Urol 2002;168:19–22.

[34] Müller V, Miszczuk M, Althoff C, Stroux A, Greiner A, Kuivaniemi H, et al. Comorbidities Associated with Large Abdominal Aortic Aneurysms Aorta 2019;in press.

[35] Bosniak MA. The current radiological approach to renal cysts. Radiology 1986;158:1–10.

[36] Alnassar S, Bawahab M, Abdoh A, Guzman R, Al Tuwaijiri T, Louridas G. Incisional hernia postrepair of abdominal aortic occlusive and aneurysmal disease: five-year incidence. Vascular 2012;20:273–7.

[37] Barisione C, Garibaldi S, Brunelli C, Balbi M, Spallarossa P, Canepa M, et al. Prevalent cardiac, renal and cardiorenal damage in patients with advanced abdominal aortic aneurysms. Intern Emerg Med 2015.

[38] Chun KC, Teng KY, Chavez LA, Van Spyk EN, Samadzadeh KM, Carson JG, et al. Risk factors associated with the diagnosis of abdominal aortic aneurysm in patients screened at a regional Veterans Affairs health care system. Ann Vasc Surg 2014;28:87–92.

[39] Takeuchi H, Okuyama M, Uchida HA, Kakio Y, Umebayashi R, Okuyama Y, et al. Chronic Kidney Disease Is Positively and Diabetes Mellitus Is Negatively Associated with Abdominal Aortic Aneurysm. PLoS One 2016;11:e0164015.

[40] Sinden NJ, Baker MJ, Smith DJ, Kreft JU, Dafforn TR, Stockley RA. Alpha-1-antitrypsin variants and the proteinase/antiproteinase imbalance in chronic obstructive pulmonary disease. Am J Physiol Lung Cell Mol Physiol 2015;308:L179–90.

[41] Song BG, Park YH. Presence of renal simple cysts is associated with increased risk of abdominal aortic aneurysms. Angiology 2014.

[42] Spanos K, Rountas C, Saleptsis V, Athanasoulas A, Fezoulidis I, Giannoukas AD. The association of simple renal cysts with abdominal aortic aneurysms and their impact on renal function after endovascular aneurysm repair. Vascular 2016;24:150–6.

[43] Brownstein AJ, Bin Mahmood SU, Saeyeldin A, Velasquez Mejia C, Zafar MA, Li Y, et al. Simple renal cysts and bovine aortic arch: markers for aortic disease. Open Heart 2019;6:e000862.

[44] Kim SM, Chung TH, Oh MS, Kwon SG, Bae SJ. Relationship of simple renal cyst to hypertension. Korean J Fam Med 2014;35:237–42.

[45] Terada N, Ichioka K, Matsuta Y, Okubo K, Yoshimura K, Arai Y. The natural history of simple renal cysts. J Urol 2002;167:21–3.

[46] Kurata A, Inoue S, Ohno S, Nakatsubo R, Takahashi K, Ito T, et al. Correlation between number of renal cysts and aortic circumferences measured using autopsy material. Pathol Res Pract 2013;209:441–7.

[47] Torres VE, Wilson DM, Hattery RR, Segura JW. Renal stone disease in autosomal dominant polycystic kidney disease. Am J Kidney Dis 1993;22:513–9.

[48] Reeps C, Maier A, Pelisek J, Hartl F, Grabher-Meier V, Wall WA, et al. Measuring and modeling patient-specific distributions of material properties in abdominal aortic aneurysm wall. Biomech Model Mechanobiol 2013;12:717–33.

[49] Mehta M, Veith FJ, Lipsitz EC, Ohki T, Russwurm G, Cayne NS, et al. Is elevated creatinine level a contraindication to endovascular aneurysm repair? J Vasc Surg 2004;39:118–23.

